# Self-organization of Tissue Growth by Interfacial Mechanical Interactions in Multi-layered Systems

**DOI:** 10.1101/2021.03.04.433890

**Authors:** Tailin Chen, Yan Zhao, Xinbin Zhao, Shukai Li, Jialing Cao, Jing Du, Yanping Cao, Yubo Fan

**Affiliations:** Key Laboratory for Biomechanics and Mechanobiology of Chinese Education Ministry, School of Biological Science and Medical Engineering, Beihang University, Beijing 100191, China; Institute of Biomechanics and Medical Engineering, Department of Mechanical Engineering, School of Aerospace, Tsinghua University, Beijing 100084, China; State Key Laboratory of Advanced Design and Manufacturing for Vehicle Body, College of Mechanical and Vehicle Engineering, Hunan University, Changsha 410082, China; Beijing Advanced Innovation Centre for Biomedical Engineering, Beihang University, Beijing 100191, China; Key Laboratory of Human Motion Analysis and Rehabilitation Technology of the Ministry of Civil Affairs, National Research Center for Rehabilitation Technical Aids, Beijing 100176, China

**Keywords:** compression gradient, interfacial interaction, morphogenesis, biomechanics, self-organization, tissue fluidity

## Abstract

Morphogenesis is a spatially and temporally regulated process involved in various physiological and pathological transformations. In addition to the associated biochemical factors, the physical regulation of morphogenesis has attracted increasing attention. However, the driving force of morphogenesis initiation remains elusive. Here, we show that during the growth of multi-layered tissues, morphogenetic process can be self-organized by the progression of compression gradient stemmed from the interfacial mechanical interactions between layers. In tissues with low fluidity, the compression gradient is progressively strengthened during growth and induces stratification by triggering symmetric-to-asymmetric cell division reorientation at the critical tissue size. In tissues with high fluidity, compression gradient is dynamic and induces cell junction remodelling regulated cell rearrangement leading to 2D in-plane morphogenesis instead of 3D deformation. Morphogenesis can be tuned by manipulating tissue fluidity, cell adhesion forces and mechanical properties to influence the progression of compression gradient during the development of cultured cell sheets and chicken embryos. Together, the dynamics of compression gradient arised from interfacial mechanical interaction provides a conserved mechanism underlying morphogenesis initiation and size control during tissue growth.

## Introduction

Morphogenesis is a common process that occurs widely in embryonic development, tissue regeneration and cancer progression. The orchestration of morphogenetic processes is complex and involves spatial and temporal regulation by biochemical factors (e.g., cell polarity signals and morphogen gradient) and physical factors (Gallet, 2011; Heisenberg and Bellaiche, 2013; Nishimura and Takeichi, 2008). Emerging studies have revealed that proper morphogenesis relies on the mechanical force of the cells and their environment. For example, apical constriction of cells caused by the contractility of myosin (Martin et al., 2009), cell junction remodelling (e.g., cell intercalation) (Rauzi et al., 2010) and tissue stiffness-dependent cell migration (Barriga et al., 2018) have been reported as important mechanisms in tissue shaping during embryonic development. These studies indicate the essential functions of cellular and molecular mechanics in the progression of morphogenesis. However, the upstream events, especially the initial driving forces of morphogenesis, remain unknown.

Most biological tissues have multi-layered structures. The interactions between layers are essential in tissue homeostasis maintenance and morphogenesis during embryonic development and pathological progression (Bhowmick and Moses, 2005; Carvalho and Heisenberg, 2010; Lilly, 2014). For example, the interaction between cancer cells and their adjacent stroma plays a key role in the progression of the diseases, including tumour invasion (Gupta and Massague, 2006; Mueller and Fusenig, 2004). In addition to biochemical communications, increasing evidence has shown the essential role of physical interactions between adjacent layers in the regulation of morphogenesis (Bailles et al., 2019; Barriga et al., 2018; Carvalho et al., 2009; Munster et al., 2019). For instance, follicle formation in chicken embryos is initiated by mechanical forces transduced from the dermal layer to the epidermal layer (Shyer et al., 2017). The villi of human and chicken guts are formed by the compressive stresses generated by smooth muscle layers on the endoderm and mesenchyme layers (Shyer et al., 2013). These studies reveal that morphogenesis processes are dependent on the mutual collaboration and mechanical compatibility of multiple layers. In this sense, it is necessary and important to address the general mechanism underlying the initiation of morphogenesis during the growth of various multi-layered tissues.

Here, by combining *in vitro* and *in vivo* biological experiments, theoretical analysis and numerical simulations, we report a general mechanism underlying the initiation of morphogenesis driven by the progression of compression gradient stemmed from the interfacial mechanical interactions between growing tissue layers.

## Results

### Progressive compression gradient is strengthened in epidermal layer during chicken feather follicle morphogenesis

During the development of avian skin, the feather follicles develop by the stratification of single-layered epidermis (Mayerson and Fallon, 1985). In the *in vitro* culture of chicken skins, we found that the feather primordia emerged at Day 2 in cultured epidermis combined with dermis after isolation from HH30 stage embryos (Figure 1a-c). However, when epidermal cell sheet was isolated and cultured alone (without dermal cell layer), it failed to stratify and no primordium was formed (Figure 1b and c), indicating that the interaction between epidermal and dermal layers is essential for the morphogenetic process of epidermal cell sheet. Moreover, during the evolution of epidermis from monolayer to multilayer, the shapes of epidermal cells showed significant alteration from flat to columnar and correlated with the stages in embryo development and with the increased epidermal cell layer number (Figure 1d and e). The deformation degree of epidermal cells was gradually declined with the increased distance to the primordium center (Figure 1f and g). These experimental observations indicate that in the beginning of follicle morphogenesis, a local compression gradient was progressively strengthened in the epidermal cell sheet accompanied with the 3D deformation of epidermis. According to our previous studies about the surface wrinkling pattern formation in a non-living chemical film/substrate composite soft material, the progression of compression gradient could be generated by mismatch deformation between adjacent layers through interfacial mechanical interactions, which leads to intriguing morphogenesis (Han et al., 2015; Zhao et al., 2015a) (Figure S1). Indeed, it has been reported that during the development of chicken skins, dermal cells form aggregations and compress the adjacent epidermal cells to form follicle primordium (Ho et al., 2019; Shyer et al., 2017). Thus, we hypothesize that the mismatch deformation between epidermal and dermal layers by dermal cell aggregation generates compression gradient through interfacial mechanical interactions and causes stratification of epidermis. However, whether tissue stratification could be triggered by the mechanical interactions between adjacent layers during tissue growth remains elusive.

**Figure 1.**
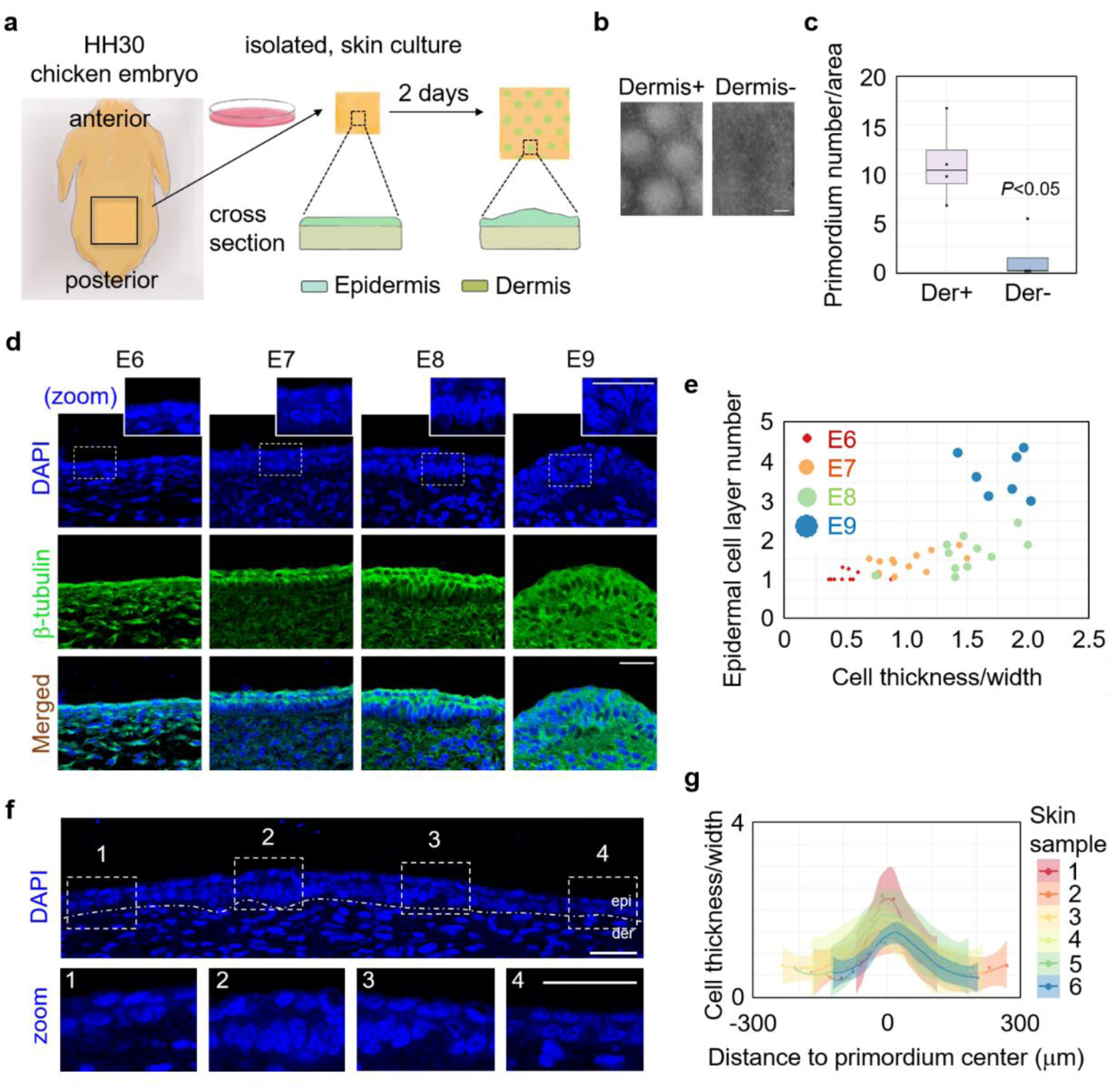
Progressive compression gradient is strengthened in epidermal layer during chicken feather follicle morphogenesis. (a) The illustration of experimental workflow for studying the epidermal morphogenesis in cultured chicken skins. (b) *In vitro* culture of embryonic chicken skin with (Der+) or without (Der-) dermal cell layer. Scale bar: 500 μm. (c) The statistical analysis of primordium number per area (mm^2^) in (b). (d) Images of skin tissues from different stages of chicken embryos showing the deformation of epidermal cell shape. Scale bar: 25 μm. (e) The statistical analysis of cell deformation (thickness/width) and cell layer number of epidermis in skin tissues from different stages of chicken embryos. Each dot represents the average value of an embryo. (f) Images of epidermal cell shape deformation at different location around the center of primordium. Scale bar: 25 μm. (g) The statistical analysis of cell deformation (thickness/width) with different distance to the center of primordium.

### 3D tissue morphogenesis could be self-organized at a critical size in multi-layered system

To study the role of interfacial mechanical interactions between layers in the initiation of tissue morphogenesis, we developed a simple film/substrate system composed of a freely growing monoclonal cell sheet and extracellular matrix (ECM) (Figure 2a). To ensure the occurrence of stratification, cells without contact inhibitory properties were studied. During the continuous live imaging, the emergence of a 2D monolayer-to-3D multilayer transition was stably observed at a critical tissue size in a wide variety of cell types including skin-derived cells (B16F10). The critical size was relatively constant for a given cell type, indicating a self-organized mechanism of morphogenesis during cell sheet growth (Figure 2a-e). In addition, similar phenomenon was observed in different types of ECM as well as altered substrate stiffness (Table S1). The behaviours of individual cells during cell sheet growth prior to the 3D morphogenesis transition were examined using a holographic imaging cytometer. Although the cell proliferation rate was negligibly altered during cell sheet expansion (Supplementary Figure S2a), the average area of individual cells significantly decreased, and the average cell thickness concurrently increased during cell sheet growth, indicating significant cell deformation (Figure 2f-h). Moreover, the largest cell deformation was observed at the central region of the cell sheet (Figure 2i-l). Single cell tracing of cell deformation also suggested significant compression of cells in the central region of cell sheet (Supplementary Figure S3). This cell deformation behaviour indicates that during cell sheet growth, a compression gradient within the cell sheet emerges. To further verify the compression gradient, a scratching experiment was performed crossing the center to the edge of the cell sheet. After scratching, cells in the central region showed much faster expansion and migration speed compared with peripheral cells, indicating the release of compressive strain (Figure S4). Similar mechanical gradients have also been obtained by Traction Force Microscope (TFM) and Monolayer Stress Microscopy (MSM) in previous studies (Perez-Gonzalez et al., 2019; Puliafito et al., 2012; Trepat et al., 2009).

**Figure 2.**
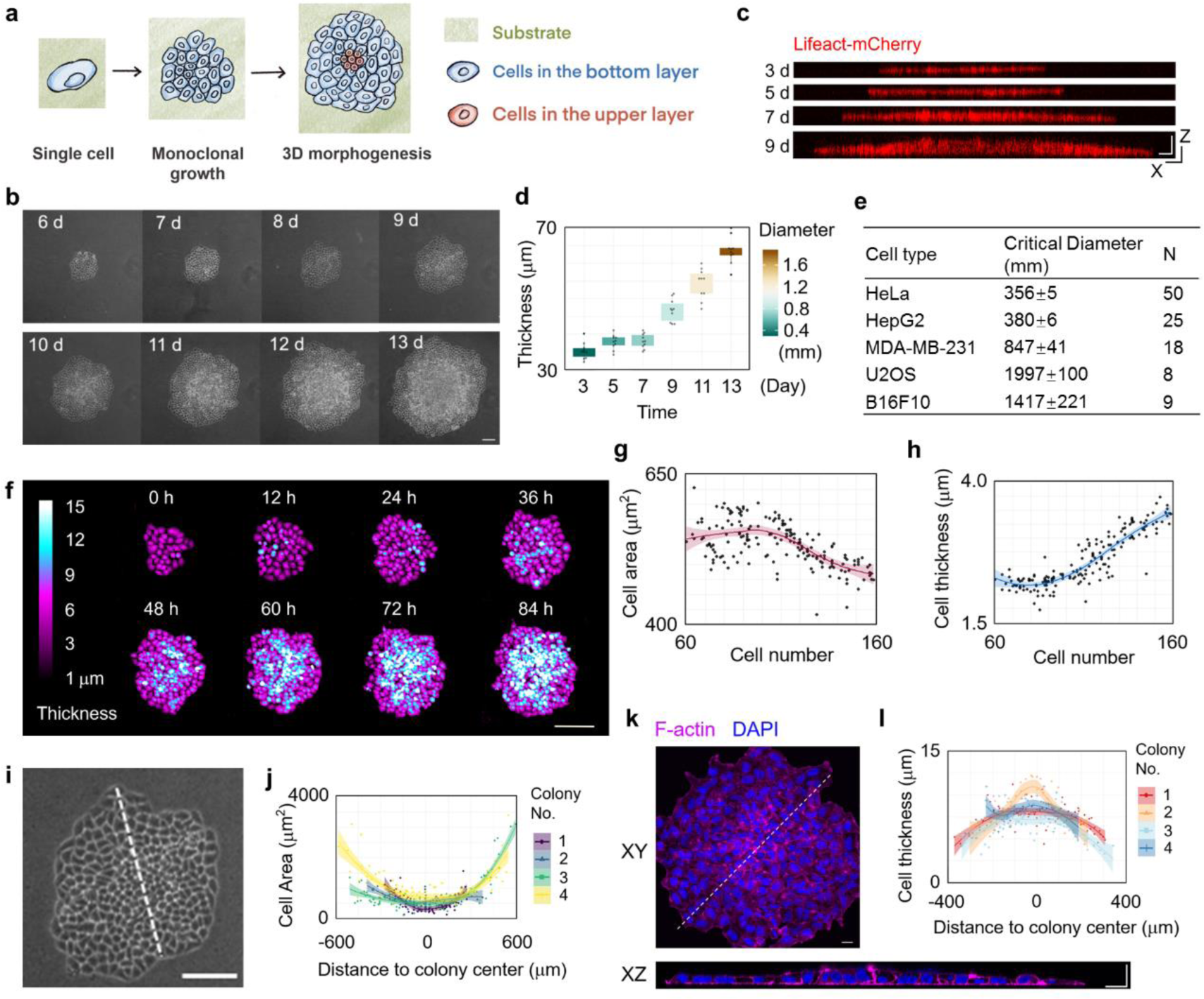
Emergent 3D morphogenesis in a growing monoclonal cell sheet at critical size. (a) The illustration of experimental workflow for studying the emergent 3D morphogenesis in a freely growing monolayer HeLa cell sheet. (b) Representative phase contrast images of a growing monoclonal HeLa cell sheet captured at the indicated time (d: day) after seeding. Scale bar: 100 μm. (c-d) XZ slice images of F-actin indicated by Lifeact-mCherry in a growing monoclonal Lifeact-mCherry^+^ HeLa cell sheet captured at the indicated time after seeding. Scale bar: 50 μm. The statistical analysis of cell sheet thickness and diameter of a growing HeLa cell sheet. (n = 10). (e) The critical size for 3D morphogenesis of growing cell sheet in different cell types. (f) The representative live images of a growing HeLa cell sheet using HoloMonitor M4 time-lapse cytometer. Scale bar: 150 μm. (g) The statistical analysis of the area of individual cell during HeLa cell sheet growth. (h) The statistical analysis of the thickness of individual cell during HeLa cell sheet growth. (i) The magnified view of HeLa cell sheet at 8 d in (b). Scale bar: 100 μm. (j) The statistical analysis of the individual cell area along the lines in (i). (k) Representative XY and XZ slice images of F-actin and nucleus stained by Phalloidin and DAPI respectively in HeLa cell sheet. Scale bar: 20 μm. (l) The statistical analysis of the individual cell thickness along the lines in (i).

To investigate the generation mechanism of compression gradient in the cell sheet during growth, we performed theoretical analysis. As illustrated in Figure 3a, the cell sheet is considered as continuum material. During cell sheet expansion caused by cell proliferation, interfacial shear stress (ISS) would be generated between the cell sheet and substrate layers. Since the direction of ISS is contrary to the relative motion between the adjacent layers, the cell sheet is subjected to ISS directed toward the center, which is consistent with the previous observations by TFM (Trepat et al., 2009). Thus, cells in the central region of the cell sheet would sustain higher level of compression than those in other regions (Figure 3a and Supplementary Mechanical Modelling), which is confirmed by the experimental observations (Figure 2i-l). The elastic strain energy stored in the cell would increase with the expansion of cell sheet. When the elastic strain energy in the cell is small, interfacial normal adhesion would impose restriction on delamination, making the cell monolayer grow in plane. Thus, higher compressive strain would be generated further. When the compressive strain reaches a critical value, elastic strain energy stored in the cell may be greater than the energy of interfacial normal adhesion and led to the occurrence of interfacial delamination. In this critical condition, stratification may happen, and the elastic strain energy can be released (Supplementary Mechanical Modelling). Indeed, we found that when cell sheet grew beyond the critical size, amounts of cells in the central region were delaminated from the substrate (Figure 3b). Theoretical analysis can also give valid quantitative predictions of experiments. First, theoretical results of the distribution of cell areas agree well with the experimental results during cell sheet expansion (Figure 3b). According to the theoretical analysis, the critical size of the cell sheet for 3D morphogenesis depends on the mechanical properties of cell sheet and substrate and interactions between them. This is confirmed by finite element simulations which well resembled the morphogenesis at the cell sheet center and also indicated that, when the compressive strain exceeded the critical value, alterations in the cell-substrate interactions could significantly affect the compression gradient during cell sheet growth (Figure 3d and Supplementary Mechanical Modelling). Moreover, experimentally manipulating cell-substrate interactions using the integrin inhibitor RGD significantly attenuated the compression gradient and morphogenesis (Figure 3e-g). In addition, inhibition of the contractility of the cell sheet using the cytoskeleton inhibitor blebbistatin or myosin shRNA significantly disrupted the compression gradient and morphogenesis (Figure 3e-g and Supplementary Figure S5). Based on the theoretical analysis, adhesion between cells may also affect the compression gradient. Consistently, inhibition of the cell-cell adhesion force by an E-cadherin neutralizing antibody also reduced the maximum compressive strain in the cell monolayer and inhibited the morphogenetic process (Figure 3e-g). These results show that the emergent 3D morphogenesis of a cell sheet/substrate system could be physically triggered by interfacial mechanical interactions between adjacent layers during growth.

**Figure 3.**
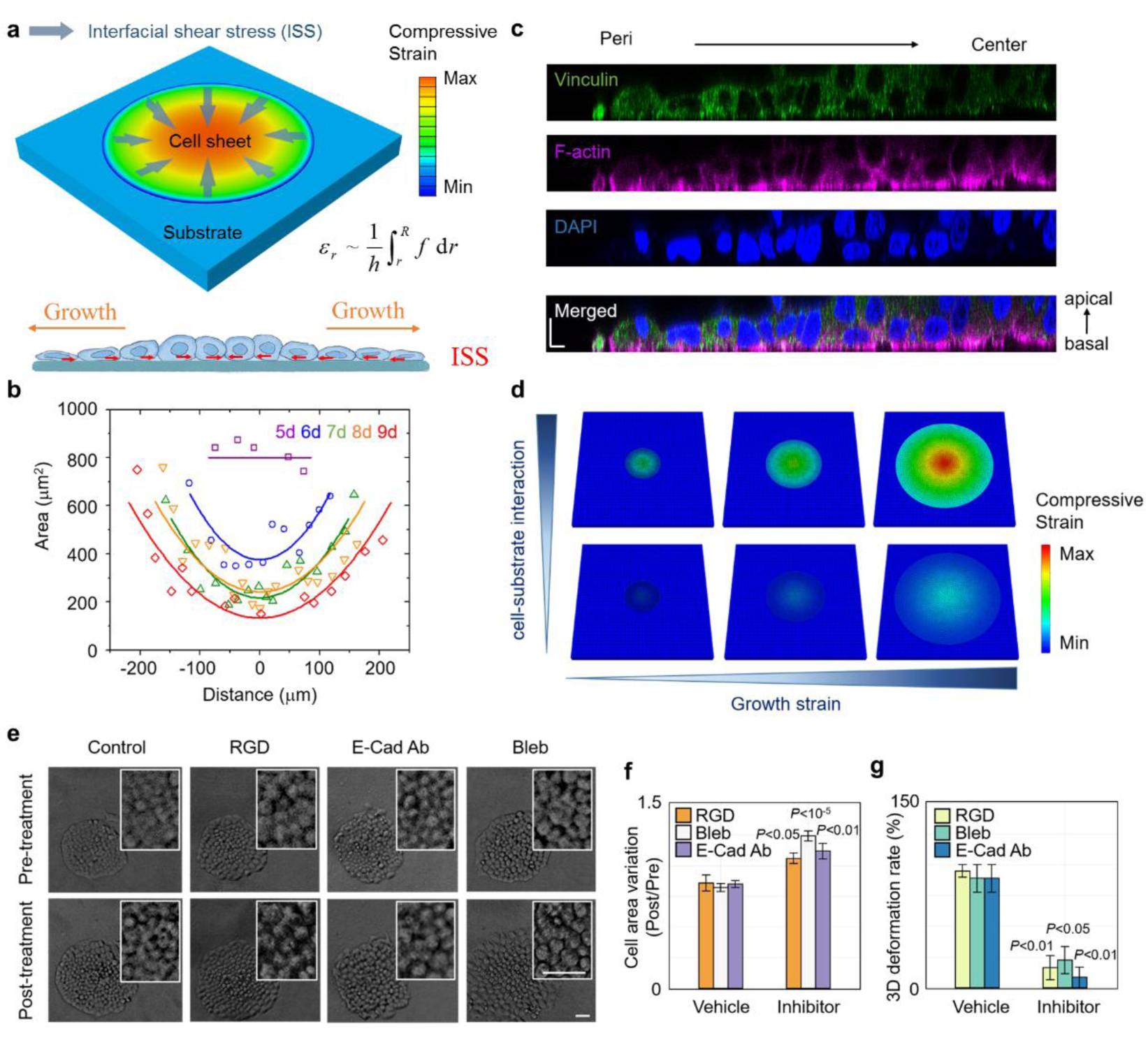
Size-dependent 3D morphogenesis is induced by interfacial shear stress in multi-layered system. (a) Theoretical model of a cell monolayer sheet growing on a substrate. *f* is the interfacial shear stress between the cell sheet and substrate generated by relative motion. *h* is the thickness of cell sheet. *σ_r_* is the compressive stress in the cell sheet induced by the interfacial shear stress. *R* is the outer radius of the cell sheet and *r* is the distance between cell and center of the monolayer sheet. Diagrams of interfacial shear stress between cell and substrate. Direction of interfacial shear stress is contrary to the cell sheet expansion, as the yellow arrow points out. (b) Area of the individual cell along a random line across the center of the cell sheet. Data points refer to the experimental results and corresponding lines are theoretical predictions obtained by fitting the experimental data using Eq. 6 in Supplementary Materials. (c) Representative XZ slice image of Vinculin, F-actin and nucleus stained by Vinculin antibody, Phalloidin and DAPI respectively in half of a HeLa cell sheet. Scale bar: 10 μm. (d) Finite element simulation of the stress field in the growing cell monolayer with different cell-substrate interactions (*f*_top_ / *f*_bottom_ = 5). Growth strains from left to right are 10%, 60% and 120%, respectively. Color bar shows the distribution of the compressive stress in the cell monolayer. (e) Representative phase contrast images of HeLa cell sheet in the presence of inhibitors or vehicle. Scale bar: 50 μm. (f) The statistical analysis of the individual cell area of HeLa cell sheet in the presence of inhibitors or vehicle. (n = 10). (g) The morphogenesis percentage of HeLa cell sheet in the presence of inhibitors or vehicle. (n = 3).

### Critical Compression triggers symmetric-to-asymmetric reorientation of cell division

We proceeded to investigate the biological mechanism of the cell layer number increase in the central region of the cell sheet induced by cell layer/substrate interfacial interaction. We found that, in contrast to symmetric (parallel) cell division in the peripheral region of large cell colonies, abundant asymmetric (oblique or perpendicular) cell division was observed in the central cells which bore significant compressive strains (Figure 4a-f). The orientation of the division plane was closely correlated with cell thickness (Figure 4f). Moreover, after release of compressive strain by cell scratching, the reorientation of cell division was diminished (Figure S6), indicating the effect of compression gradient on cell division orientation. This asymmetric cell division ultimately led at least one daughter cell to locate at the top layer of the cell sheet (Figure 4g and Supplementary Movies 1 and 2). The cell division reorientation induced by compression gradient was also confirmed by the subcellular localization of the nuclear-mitotic apparatus protein (NuMA), which is involved in the orchestration of mitotic spindle positioning (Pirovano et al., 2019) (Figure 4h). Moreover, spindle-rocking experiments indicate that cells in the central region showed a significantly higher level of metaphase plate oscillations which is always associated with asymmetric cell division (Haydar et al., 2003) (Figure 4i). These results suggest that compression gradient induced by interfacial interaction between layers triggers cell division reorientation from symmetric to asymmetric leading to tissue stratification.

**Figure 4.**
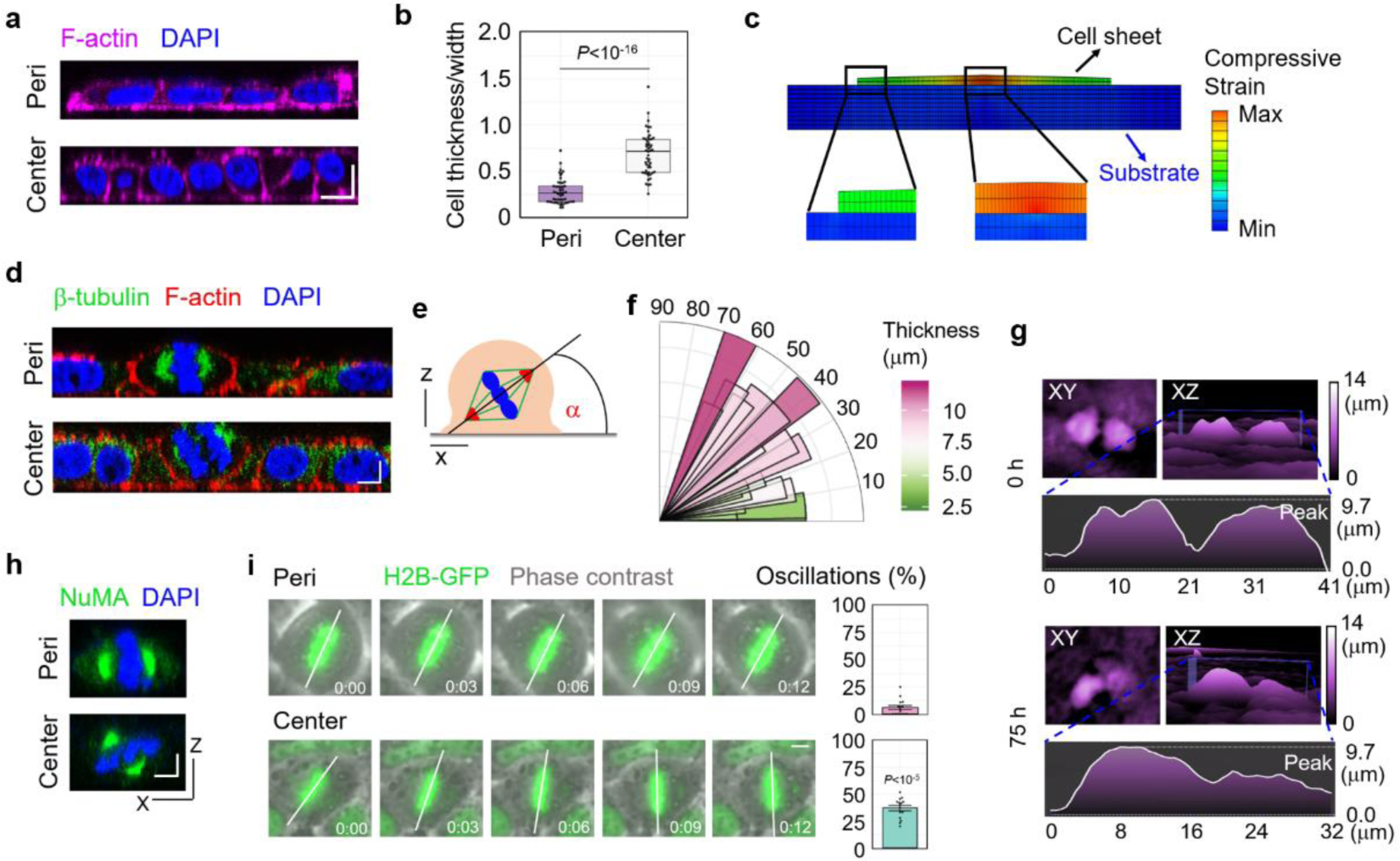
Critical Compression triggers cell division reorientation to induce tissue stratification. (a) Representative XZ slice image of F-actin and nucleus stained by Phalloidin and DAPI respectively in the central region and peripheral region of HeLa cell sheet. Scale bar: 10 μm. (b) The statistical analysis of cell deformation (thickness/width) in the center region and peripheral region of Hela cell sheet. (n = 50). (c) Cell shape variation induced by interfacial shear stress. (d) Representative XZ slice image of β-tubulin, F-actin and nucleus stained by β-tubulin antibody, Phalloidin and DAPI respectively in the central region and peripheral region of HeLa cell sheet. Scale bar: 5 μm. (e) The schematic experimental setting of the mitotic spindle orientation. (f) Distribution of the spindle-axis angles of cells with different thickness in HeLa cell sheet. (g) The representative images of dividing cells before (0 h) and after (75 h) critical compression during HeLa cell sheet growth using HoloMonitor M4 time-lapse cytometer. (h) Representative XZ slice image of NuMA and nucleus stained by NuMA antibody and DAPI respectively in the central region and peripheral region of HeLa cell sheet. Scale bar: 5 μm. (i) Analysis of spindle oscillation in in the central region and peripheral region of GFP-H2B^+^ U2OS cell sheet. The extent of oscillation was calculated and plotted in bar graphs on the right. Scale bar: 5 μm. (n = 14).

### Tissue fluidity controls 2D/3D morphogenetic transition by regulating compression gradient progression

Our experimental results show large variations in the critical sizes of cell sheets for 3D morphogenesis among different cell types with critical diameter ranging from 356 mm in HeLa to 1997 mm in U2OS cells (Figure 2e). Notably, some cell types, such as Madin-Darby canine kidney (MDCK) cells, did not display stratification during our observations. To further investigate the intrinsic properties of tissue layers that affect the emergent morphogenesis, we compared cell behaviours during cell sheet growth between HeLa and MDCK cells. First, in order to exclude the influence of contact inhibition on MDCK cell behaviour, we measured cell proliferation rate and cell density alterations during cell sheet growth. As shown in Supplemental Figure S2b and d, MDCK cells did not show contact inhibition during our observation. Moreover, according to the previous report that MDCK cell sheet did not display contact inhibition until a critical size of approximately 2×10^6^ μm^2^ (Puliafito et al., 2012). In our experiment, the cell sheet size of MDCK colony was below 1×10^5^ μm^2^. Thus, the differential behavior of MDCK cell sheet was not caused by contact inhibition. Interestingly, we found that, instead of 3D morphogenesis, MDCK cell sheet showed dramatic alterations in its in-plane shape during growth, with dynamically organized protrusions at cell sheet edges, indicating an emergent 2D morphogenesis (Supplementary Figure S7). While few differences in the proliferation rate were observed between these two cell types (Supplementary Figure S2a and b), expansion rate was significantly higher in MDCK cell sheet than in HeLa cell sheet during growth (Supplementary Figure S2c and d). Importantly, in contrast with the HeLa cell sheet behaviour, the MDCK cell sheet did not show progressive compression gradient strengthen during growth (Figure 5). Moreover, while the individual cell area in the HeLa cell sheet was relatively steady, MDCK cells exhibited strong area fluctuation during cell sheet growth (Figure 6a and b). Particle imaging velocimetry (PIV) analysis showed that during cell sheet expansion, MDCK cells displayed rapid collective cellular motion with a higher overall cell velocity (root-mean-square, rms velocity, v_rms_), indicating a fluid-like state (Garcia et al., 2015). In contrast, the collective behaviour of HeLa cells showed slower cell motion, indicating a relatively solid-like state (Figure 6c, d and Supplementary Movie 3). The fluidity of these two cell types was further confirmed by the cell shape index p_0_, the median ratio of the perimeter to the square root area of the cells. According to the vertex model, if the cell shape index of the system increases to p*_0_ ≈ 3.81, a transition from a jammed, solid-like state to an unjammed, fluid-like state occurs (Bi et al., 2015). Over the course of cell sheet growth, the average cell shape index of MDCK cells was constantly above 3.81, whereas HeLa cells approached the jamming threshold p*_0_ (Figure 6e and f). Moreover, the cell shape index was lower in the central region than in the peripheral region of the HeLa cell sheet (Supplementary Figure S8). At the edges of MDCK cell sheet, especially during the formation of protrusions, significant collective cell migration (Figure 5a and Supplementary Movie 4) and abundant cell intercalations such as T1 transitions and rosettes formation were observed (Figure 7a and b). Cell intercalation is reported as a mechanism for driving tissue extension in embryonic development (Guillot and Lecuit, 2013) and could be controlled by external constraints acting on the tissue (Aigouy et al., 2010). In our experiment, most of cell intercalation was observed in cells with relatively smaller areas (less than 300 μm^2^), indicating a high level of compressive strain (Figure 7c). Moreover, the cell junction remodelling during intercalation process was closely correlated with the 2D deformation of cell sheet, indicated by the small angles between new junctions and tissue protrusion directions (Figure 7d) and higher tissue elongation rate at the direction of new junctions (Figure 7e). To investigate the effect of the tissue fluidity on the 2D/3D morphogenetic transition, cellular migration ability was inhibited using the small GTPase Rac1 inhibitor NSC23766 (Raftopoulou and Hall, 2004). The results showed that Rac1 inhibitor treatment significantly reduced tissue fluidity (Figure 7f and g and Supplementary Movie 5) and induced higher level of compressive strain in MDCK cell sheet (Figure 7h and i). Importantly, treatment of NSC23766 significantly promoted the emergence of tissue stratification (Figure 7j). In comparison, inhibition of Rho-associated, coiled-coil containing protein kinase (ROCK) by Y27632 slightly enhanced tissue fluidity of MDCK cells (Figure 7f and g and Supplementary Movie 5) and had little effect on morphogenesis (Figure 7h-j). These results suggest that the morphogenesis driven by interfacial mechanical interactions between tissue layers is dependent on tissue fluidity that higher level of tissue fluidity may prevent the storage of the compressive strain energy through in-plane cell motion.

**Figure 5.**
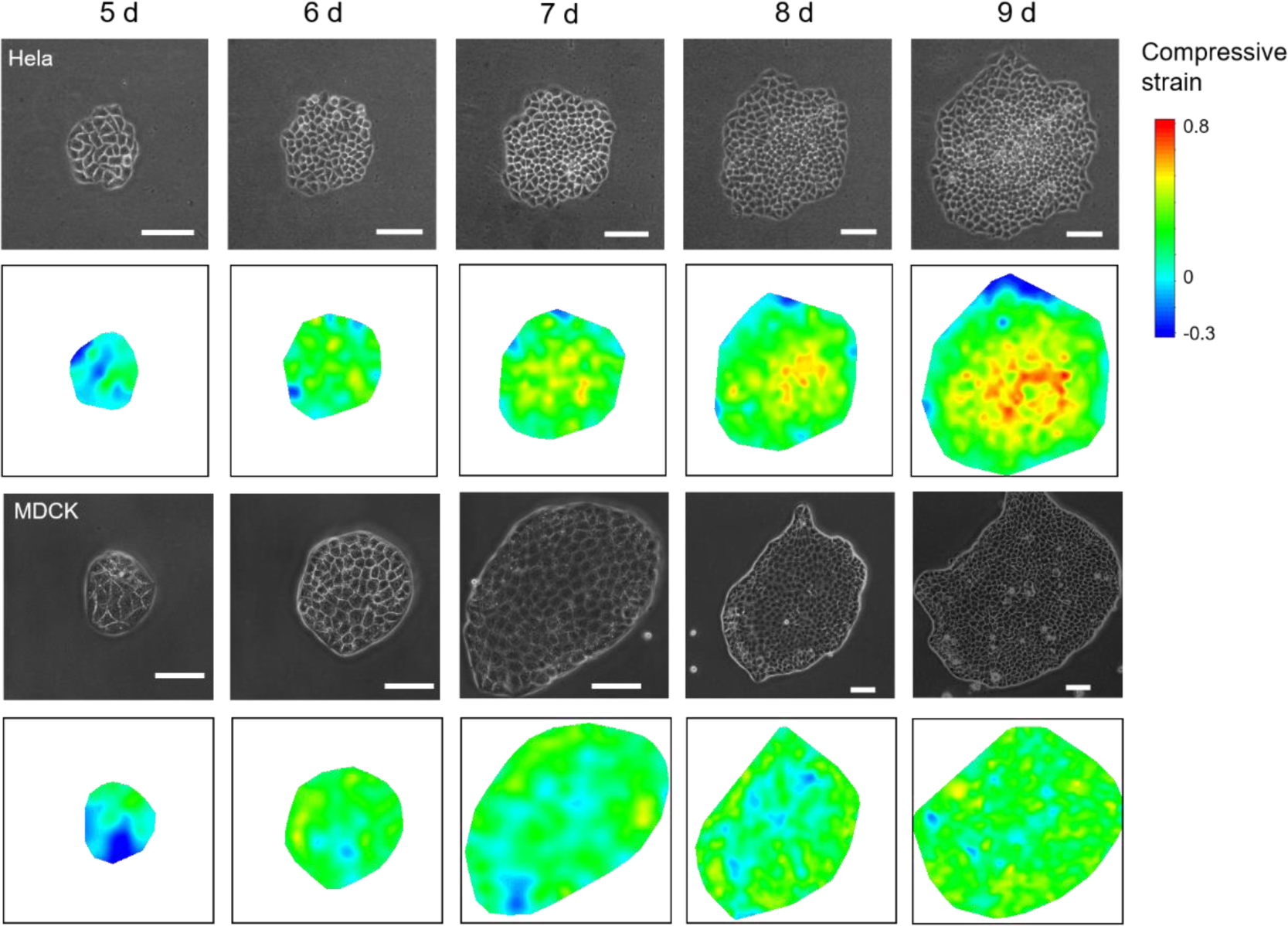
Strain field evolution of HeLa and MDCK cell sheets. The calculated strain field evolution during the growth of Hela and MDCK cell sheets at the indicated time (d: day) after seeding. Compressive strain field was calculated using Eq. (4) in the Supplementary materials by measuring the areas of each cell. The normal area of a full-grown cell without sustaining compression was set as the average area of the cell sheet in 1 d since the compressive gradient is small in the initial stage. Scale bar: 100 μm.

**Figure 6.**
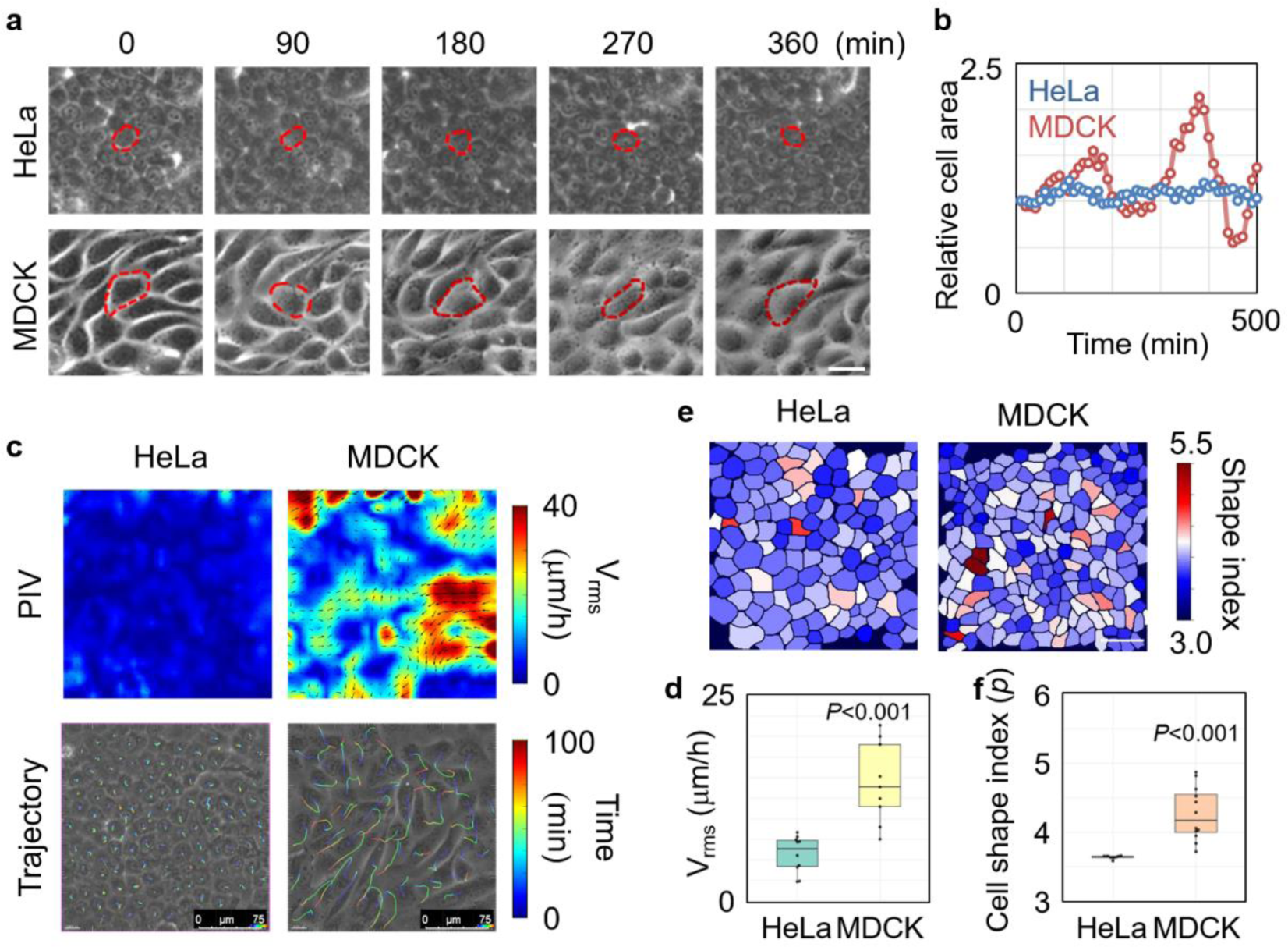
HeLa and MDCK cell sheets show different fluidity during growth. (a) The fluctuation of individual cell shape during the growth of HeLa and MDCK cell sheets. Scale bar: 25 μm. (b) The area alteration with time of the cell indicated by dotted line in (a). (c) Cell velocity field analyzed by PIV and cell trajectories in HeLa and MDCK cell sheets. Scale bar: 75 μm. (d) The statistical analysis of cell speed (rms velocity) measured by PIV in HeLa and MDCK cell sheets. (n = 10). (e) The cell shape index distribution of HeLa and MDCK cell sheets. Scale bar: 50 μm. (f) The statistical analysis of cell shape index of HeLa and MDCK cell sheets. (n = 10).

**Figure 7.**
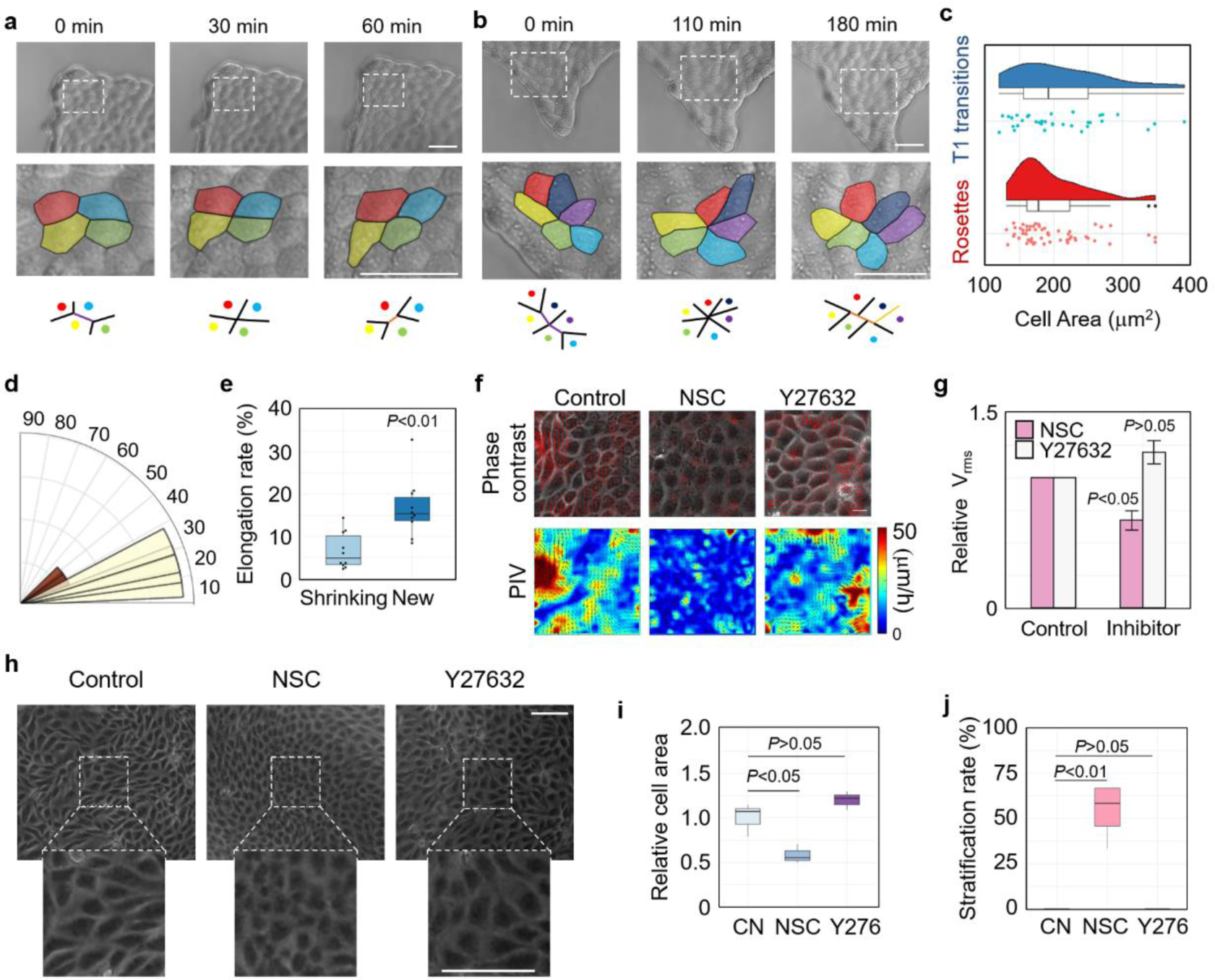
Tissue fluidity controls 2D/3D morphogenetic transition by regulating compression gradient progression. (a) Representative image of T1 transition at MDCK cell sheet edges during growth. Cell rearrangements are illustrated at the bottom layer (purple line: shrinking junction, orange line: new junction). (min: minute). Scale bar: 50 μm. (b) Representative image of rosettes at MDCK cell sheet edges during growth. Cell rearrangements are illustrated at the bottom layer (purple line: shrinking junction, orange line: new junction). Scale bar: 50 μm. (c) Cell area distributions of cells be experiencing T1 transitions and rosettes. (d) Distribution of the angles between new junctions and tissue protrusion directions. (e) Cell sheet elongation rate at the direction of shrinking junctions or new junctions. (n = 10). (f) The velocity field superimposed on the corresponding phase contrast (upper panel) and velocity map (lower panel) images measured by PIV of MDCK cell sheet in the presence of inhibitors or vehicle. Scale bar: 20 μm. (g) The statistical analysis of cell speed (rms velocity) measured by PIV of HeLa and MDCK cell sheets in the presence of inhibitors or vehicle. (n = 4). (h) Representative phase contrast images of MDCK cell sheet in the presence of inhibitors or vehicle. Scale bar: 100 μm. (i) The statistical analysis of the individual cell area of MDCK cell sheet in the presence of inhibitors or vehicle. (n = 3). (j) The stratification percentage of MDCK cell sheet in the presence of inhibitors or vehicle. (n = 4).

### Progression of compression gradient contributes to epidermal cell stratification during chicken skin development

Asymmetric cell division is a common mechanism involved in tissue stratification and cell fate differentiation during embryogenesis and cancer progression (Lechler and Fuchs, 2005; Neumuller and Knoblich, 2009). There is evidence that during the development of chicken skins, the deformation of basement membrane between epidermis and dermis by the aggregation of dermal cells plays crucial role for the initiation of follicle primordium formation (Ho et al., 2019; Shyer et al., 2017). According to our results, tissue stratification could be initiated by the mechanical interaction at interface between adjacent layers through compression-induced cell division reorientation. Thus, we proceeded to verify whether this mechanism is applied to the stratification of epidermis during chicken feather follicle morphogenesis. First, we compared the cell division orientation of the epidermis in single-layered stage and multi-layered stage during skin development. We found that when epidermis was stratified, most of the mitotic spindles of epidermal cells were reoriented from symmetric to asymmetric and the spindle-axis angle was closely correlated with cell deformation degree (Figure 8a-c). Moreover, reinforcing the progression of compression gradient by reducing tissue fluidity using Rac1 inhibitor significantly promoted the progression of epidermis stratification and follicle formation, indicated by the epidermal cell layer number and pattern geometry of follicle primordium (Figure 8d-i). Reducing tissue fluidity also increased the nuclear localization of β-catenin protein, indicating the promotion of follicle cell fate determination (Figure 8h and i). In addition, disrupting the interfacial mechanical interaction between epidermal and dermal layers by inhibiting epidermal cell/base membrane adhesion using integrin inhibitor largely attenuated the morphogenesis of primordium (Supplementary Figure S9). Thus, our findings suggest a model for the initiation of epidermis stratification during feather follicle formation that dermal cell aggregation induces mismatch deformation between epidermal layer and basement membrane which generates progressive compression gradient in epidermal layer and the latter triggers cell division reorientation leading to epidermis stratification.

**Figure 8.**
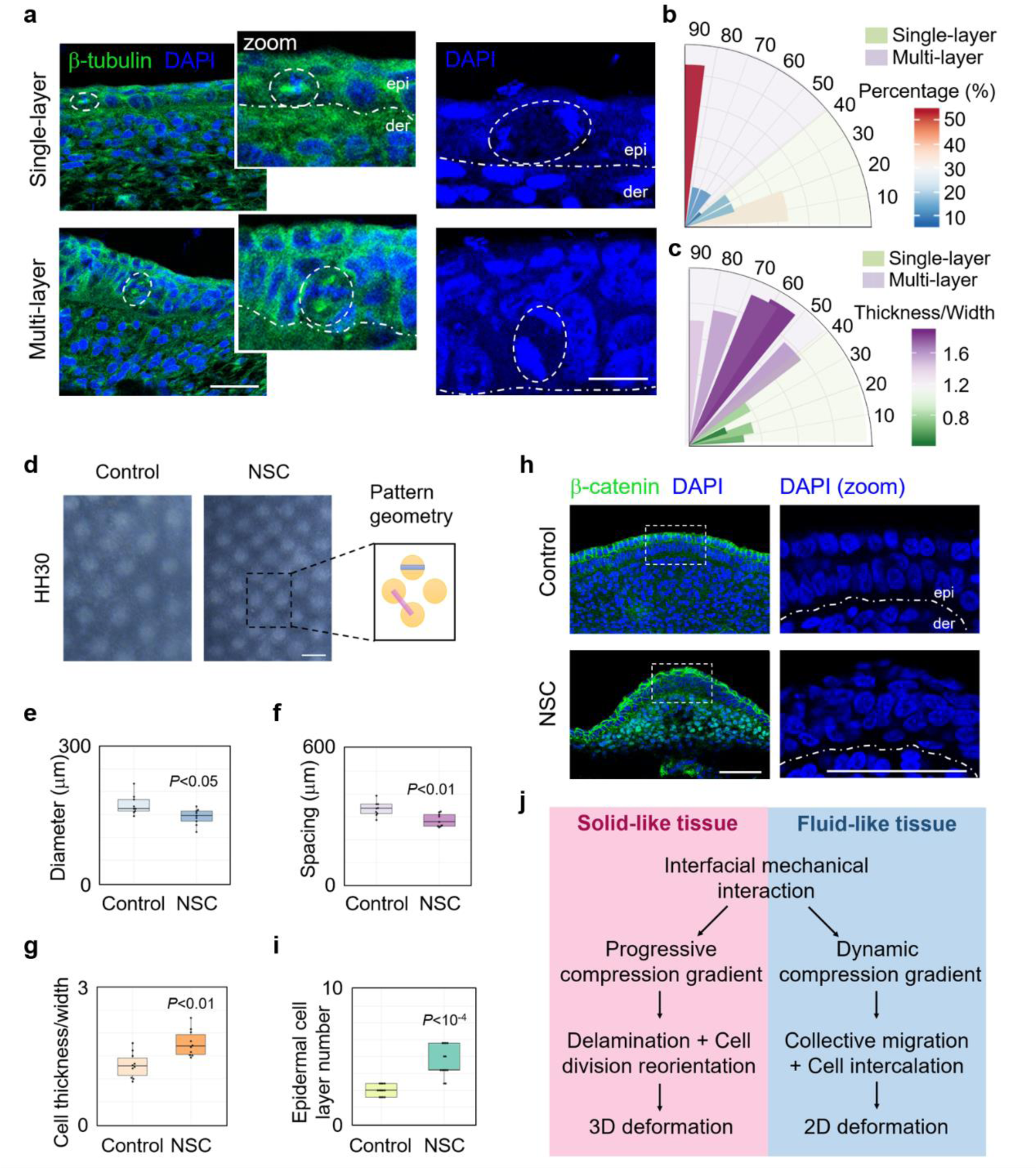
Progression of compression gradient contributes to epidermal cell stratification during chicken skin development. (a) Images of embryonic chicken skin showing the orientation of mitoses relative to the basement membrane (white dotted line) in the magnified images (zoom), separating epidermis (epi) from dermis (der). White dotted circles indicate the mitotic cells in metaphase (left) and anaphase (right). DAPI marks the DNA and β-tubulin antibody immunofluorescence staining marks the spindle. Scale bar: 10 μm. (b) Distribution of the spindle-axis angles of cells in single-layered or multi-layered epithelia of embryonic chicken skin. (c) Distribution of the spindle-axis angles of cells with different cell deformation (thickness/width) in single-layered or multi-layered epithelia of embryonic chicken skin. (d) Reconstitution culture of embryonic chicken skin with or without NSC23766 (NSC). Scale bar: 500 μm. Quantification of (e) spacing and (f) diameter as pattern geometry parameters as illustrated in the right panel of (d). (n = 10). (g) Quantification of cell deformation of epidermis in in the presence of NSC or vehicle. (n = 10). (h) Representative cross section image of β-catenin and nucleus stained by β-catenin antibody and DAPI respectively in reconstitution cultured embryonic chicken skin with or without NSC23766 (NSC). Scale bar: 50 μm. (i) Quantification of layer number of epidermis in (h). (n = 14). (j) Model of 2D and 3D morphogenesis controlled by the interfacial mechanical interactions in multi-layered systems with different tissue fluidity.

## Discussion

In this work, we propose a self-organization mechanism for the initiation of morphogenesis during tissue growth in which a progressive compression gradient caused by macroscopic interfacial mechanics drives the initiation of morphogenesis at certain developing stages in multi-layered tissues. Our experiments and theoretical analysis show that mismatch deformation between adjacent layers during tissue growth induces progression of compression gradient by interfacial interactions which triggers cell delamination and cell division reorientation in solid-like state tissues and ultimately induces 3D deformation and tissue stratification. Meanwhile, in fluid-like state tissues, the interaction between adjacent layers induced dynamic progression gradient leading to 2D tissue deformation instead of 3D deformation (Figure 8j). We also proposed the crucial role of progressive compression gradient in epidermis generated by the interaction between epidermal and dermal layers in the initiation of epidermal cell stratification during chicken skin development. Interestingly, in zebrafish embryos, the friction force between mesoderm and neurectoderm layers during cell moving has been reported to be a key determinant in the positioning of neural anlage (Smutny et al., 2017). Thus, interfacial mechanical interaction between adjacent layers may be a conserved force origin during embryonic development. Resembling to the instability process in film/substrate material system, we regard the cell sheet as a film, which is subjected to the interfacial shear stress during cell sheet expansion. Thus, the growing cell sheet will be compressed and buckled. We found that cell-cell adhesion, cell-substrate adhesion, and cell fluidity all affected the degree of cell sheet compression and tissue stratification. Interestingly, recent studies regard the process of cell aggregation and layering as the wetting problem and also reveal that inhibiting cell contraction, cell-to-cell adhesion, and cell-to-substrate adhesion have significant influences on the dewetting process (Perez-Gonzalez et al., 2019; Ravasio et al., 2015).

Morphogenetic processes always occur at certain developmental stages during tissue growth. In Drosophila wing disc development, different cell proliferation rates induce mechanical strain to shape tissues and control tissue size by regulating proliferation and division orientation (Legoff et al., 2013; Mao et al., 2013). In our experimental model, 2D and 3D morphogenetic process emerges at the critical tissue size, which is determined by the critical value of the compressive strain generated by interfacial interaction during tissue growth. This macroscopic mechanical dissipation provides a possible regulatory mechanism for the spatiotemporal control of morphogenesis and size control of growing tissues. It has been recognized that crowding is a common phenomenon in physiological and pathological processes and regulates tissue homeostasis maintenance (Eisenhoffer et al., 2012; Marinari et al., 2012). A recent study reports that during the development of zebrafish heart, proliferation-induced crowding leads to tension heterogeneity that drives cell stratification (Priya et al., 2020). According to our results, critical compressive strain has profound effects on cell behaviours, including cell delamination, cell division reorientation and in-plane cell rearrangement, which all contribute to tissue shaping, homeostasis maintenance and size control. Moreover, in our model experiment, the compression gradient is autonomously generated by tissue growth and therefore can well simulate the *in vivo* crowding conditions with the controlled crowding levels.

As an important characteristic of tissues, fluidity (jamming/unjamming state) is determined by collective cell motion and dynamically altered during tissue growth (Sadati et al., 2013). Recent studies suggest that tissue fluidity plays crucial roles in homeostasis maintenance and tumour invasion (Garcia et al., 2015; Miroshnikova et al., 2018; Park et al., 2016). Saadaoui *et al*. reported that myosin contractility-induced global tissue flow contributed to morphogenesis during avian gastrulation (Saadaoui et al., 2020). In our work, we found that tissue fluidity could affect the local accumulation of compressive strain under interfacial interaction between adjacent layers, which finally regulates tissue deformation patterns (2D or 3D). These findings may provide insights into the regulatory mechanism underlying morphogenesis by tissue fluidity.

There is no doubt that mechanical forces play essential roles in the regulation of morphogenesis. The role of cell-scale forces generated by cytoskeleton contraction and transmitted by cell adhesions has been intensively studied in regulating tissue morphogenesis (Heisenberg and Bellaiche, 2013). However, the upstream regulation mechanism of these molecular machines, especially at the tissue scale, is rarely investigated. Our results reveal that progression of compression gradient controlled by the tissue-scale interfacial shear stress has propounding effects on local cell delamination, reorientation of division plane and cell intercalation to initiate morphogenesis. Moreover, the interfacial mechanics could be stemmed from mismatch deformation during cell migration, aggregation, proliferation, *etc.* Thus, this scale-spanning mechanical loop from the cell scale to the tissue scale and then returns to the cell scale may be a fundamental self-organized mechanism during morphogenetic process in growing multi-layered tissues. Further study is needed to elucidate the molecular mechanotransduction mechanism underlying the regulation of these cell behaviours by the progression of compression gradient under interfacial mechanical interactions.

Taken together, our findings unveil a self-organized mechanism that drives the initiation of morphogenesis in growing multi-layered tissues. These results also open up a new avenue that tissue-scale forces regulate cell behaviours, which could in turn facilitate tissue self-organization.

## Materials and Methods

### Cell culture and Immunofluorescence

HeLa, HepG2, Madin-Darby canine kidney (MDCK), MDA-MB-231, U2OS, B16F10 cells were cultured in DMEM medium (containing with 4.5 g/L glucose, L-glutamine, and sodium pyruvate) supplemented with 10% FBS (Life technologies, CA, USA), and 100 IU/mg penicillin-streptomycin (Life technologies, CA, USA) and 1 % (v/v) NEAA (Life technologies, CA, USA). For the experimental treatment, we use clones of cells grown for the same amount of time. The pharmacological agents were added including Blebbistatin (sigma, 25 μM), RGD (Abcam, 50 μg/ml), E-cadherin neutralizing antibody (Biolegend, 10 μg/ml), NSC23766 (Abmole, 20 μM), Y27632 (Abmole, 20 μM) if applicable. Images were taken on the Nikon microscope. Cells were grown at 37 °C in an incubator with 5% CO_2_. Cells grown on glass bottom dishes were fixed with 4% paraformaldehyde for 10 min at room temperature. Incubate the cells with 5% BSA, in PBST (PBS+0.1% Tween 20) for 2 hours. Cells were incubated with primary antibodies at the optimal concentrations (according to the manufacturer’s instructions) at 4 °C overnight. After washing, cells were incubated for 2 hours with secondary antibodies: 488/568/633 IgG (H+L) and/or Alexa Flour 568/647 phalloidin (Invitrogen) for 1 hour. Cell nucleus were stained with DAPI (4’,6-diamidino-2-phenylindole, Invitrogen) for 10 min at room temperature. Confocal images were taken on the Leica microscope equipped with a 10x, 40x or 63x objective. Experiments were replicated at least three times.

### Live imaging

Live cell imaging was performed in Leica microscope or HoloMonitor M4, enclosed in an incubator to maintain the samples at 37 °C and 5% of CO_2_ throughout the experiments. Images were acquired every 10 min with Leica software. Spindle-rocking experiments were acquired every 3 min with Leica microscope. HoloMonitor M4 is a Quantitative phase imaging-based cell analyzer utilizing the principle of digital holographic microscopy. Live cell imaging was performed in HoloMonitor M4, enclosed in an incubator to maintain the samples at 37 °C and 5% of CO_2_ throughout the experiments. Images were acquired every 10 min with HStudio 2.7.

### PIV (Particle Image Velocimetry) measurement

PIV analysis was conducted using a custom algorithm based on the MatPIV software package for MATLAB. We used a series of live cell images of HeLa and MDCK to calculate the velocity of the cells in the cell sheet. The mean velocity was subtracted from calculated velocity fields to avoid any drift-related bias and to get the velocity fields of the cells (the net movement of the cell cluster for its drift is less than 10% and can be ignored). The heat maps of magnitude are also exported subtracting the mean velocity. From the exported text files, we measured the overall cell speed, or root-mean-square (rms) velocity vrms and calculated the average vrms of each type of cell. The correlation algorithm was coded using MATLAB in our lab. Experiments were replicated at least three times.

### Image processing, segmentation and quantification

Cell data analysis of HoloMonitor M4: Cell area, cell thickness, cell sheet area and cell sheet thickness were analyzed by the software HStudio 2.7. HStudio 2.7 can automatically segment and extract physical parameters of the cell. For more details, please refer to the official manual.

The area of cell clones and individual cells: Cell boundary labeling and areas were determined manually in Fiji.

Cell thickness, width and division angle: The collected confocal tomography images were imported into Bitplane imaris for 3D reconstruction. After reconstruction, XZ and YZ profiles were randomly selected to measure thickness, width and cell division angle manually. Data was measured on three samples, using three regions of the image. Thickness, width and division angle in chicken embryo skin: The collected confocal tomography images were imported into Bitplane imaris. Cell thickness, length, and division Angle were measured manually on software.

Pattern geometry in Chicken embryo skin: Diameter and spacing was measured manually using Fuji.

Shape index: To determine the cell boundary, we used a semi-automatic segmentation pipeline. We manually enhanced the blurry boundaries of the cells. After processing the image by binary Ostu, we produced the finally image by Median filtering. We extracted the parameters by image segmentation to obtain the area and perimeter. Cell trajectory: The cell tracking image is produced by software Bitplane imaris. Cell junction analysis: Cell boundary labeling and areas were determined manually in Fiji(Firmino et al., 2016).

### Strain field analysis

he area of each cell was calculated by Fuji software, and the compression strain of each cell was calculated according to Eq (3) in Supplementary Materials. Finally, origin software is used to draw the compression strain diagram. The strain field is obtained by analyzing variation of cell area using Eq. (4) in Supplementary Materials.

### Finite element simulations

Finite element simulations were performed using commercial software ABAQUS (2016). Details of the finite element method were present in the Supplementary Materials. In the finite element model, the growing cell monolayer sheet was placed on a stiff substrate with the interfacial fraction factor being controlled. The cell sheet was under isotropic expansion to simulate cell proliferation and growth. The cell sheet was modeled as the linear elastic material. No other boundary conditions were applied on the cell sheet.

### Skin culture and immunofluorescence

Fertilized eggs were bought from local farms. The eggs were cultured in a moist environment at 37 °C and staged according to Hamburger and Hamilton. Dorsal skin pieces were dissected from E6 embryos and spread flat on the Polycarbonate membrane (pore size: 0.4 μm, Corning, Cat.No.3413). Culture is DMEM with 2% chick serum and 10% FBS along with pharmacological agents NSC23766 (Abmole, 20 μM), RGD (Abcam, 100 μg/ml) if applicable. Skin pieces were cultured for 48 hours at 37 °C before being fixed in 4% paraformaldehyde in PBS. Images were taken on the Nikon stereoscopic microscope. Pattern geometry was calculated using Fiji to measure the spacing and diameter. Skin culture details were performed as previously described. For immunofluorescence staining, Embryos or cultured dissected skin pieces were fixed in 4% paraformaldehyde in PBS and embedded in OCT. The tissue blocks were placed in liquid nitrogen for 1 minute and frozen section. The section thickness is about 14 µm. We incubate the sections with 5% BSA, in PBST (PBS + 0.1% Tween 20) for 2 hours. The sections were incubated with primary antibodies at the optimal concentrations (according to the manufacturer’s instructions) at 4 °C overnight. After being washed with PBS, the sections were incubated for 2 hours with secondary antibodies: 488/568/633 IgG (H+L) for 1 hour. Nucleus were stained with DAPI (4’, 6-diamidino-2-phenylindole, Invitrogen) for 10 min at room temperature. Confocal images were taken on the Leica microscope equipped with a 10 x, 40 x or 63 x objective. Experiments were replicated at least three times.

### Quantification and Statistical analysis

The number of biological replicates in each experimental result was indicated in the figure legends. Data were presented as means ± SEM. Significance was determined using Student’s t test to compare the differences between two experimental groups.

## Acknowledgments

### General

We thank J. Y. Fu (China Agricultural University) for providing GFP-H2B+ U2OS cells and C. Dong (The Pennsylvania State University), Y. Liang (EV-bio) and C. Y. Xiong (Peking University) for discussions during the preparation of this paper. We also thank S. Wu (China Agricultural University) and X. Liu (Beihang University) for technical support of chicken embryonic experiments and PIV analysis, respectively.

### Funding

This work was supported by the National Key R&D Program of China (2017YFA0506500, 2016YFC1102203, and 2016YFC1101100), the National Natural Science Foundation of China (31370018, 11972206, 11902114, 11421202, 11827803, and 11902020), and Fundamental Research Funds for the Central Universities (ZG140S1971).

### Author contributions

T.L.C, Y.Z., J.D., Y.P.C, and Y.B.F. designed the study, performed and interpreted experiments. X.B.Z. and S.K.L. helped with cell experiments and statistical analysis. J.L.C helped with PIV analysis. Y.Z. and Y.P.C. did mechanical analysis. J.D., Y.P.C., and Y.B.F. conceived and supervised this project and prepared the paper.

### Competing interests

There are no competing interests.

## Supplementary Materials

### Theory model

The theoretical model is shown in Fig. 3a. The cell monolayer is modeled as a thin film, which is attached on the surface of a rigid substrate. Direction of the tangential adhesion force between the cell and substrate is contrary to the relative motion. Thus, compression would be induced when the cell monolayer is growing on the substrate. The tangential adhesion force is assumed as *f* = *f* (*r*), where *r* is the distance between the cell and the center of the monolayer. *f* is concerned with the tangential adhesion between the cell and substrate, which is related to the cell type, stiffness of substrate and the rate of cell division or growth. By analyzing the stress state of the monolayer and solving the equilibrium equation, one can obtain the distribution of the equi-biaxial compressive stress in the monolayer

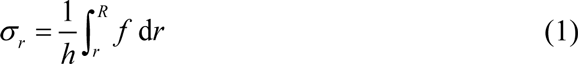

Here *R* is the outer radius of the cell monolayer, and *h* is the thickness of cell sheet. The compressive strain can be obtained by the constitutive relation as

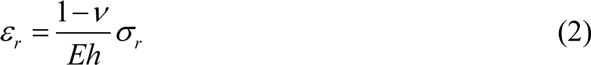

where *E,v* are the modulus and Poisson’s ratio, respectively. The central region of the monolayer would sustain higher level of compression than the cells in other regions. Thus, cell extrusion is most likely to occur in the central region, which is consistent with the experimental observations. The maximum compressive strain in the cells can be obtained as

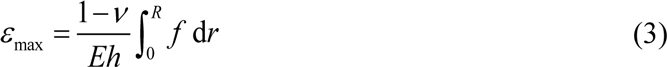

which depends on the cell-substrate interactions and area of the cell monolayer. Based on Eq. (2), one can also obtain the cell area

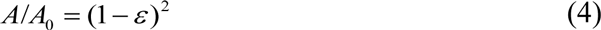

where *A*_0_ is the normal area of a full-grown cell without sustaining compression. Eq. (4) demonstrates that distribution of cell area can reflect the strain state in the cell monolayer. With cell proliferation, radius of the cell monolayer increases, and area of cells in the central region is reduced, indicating that the cell monolayer has a high level of compression in the central region.

Given the distribution of the interfacial adhesion force, one can obtain the distribution of the compressive strain and cell area of the cell monolayer. In this work, the tangential adhesion force is assumed to be uniform over the cell monolayer. Thus, the maximum compressive strain in the cell and the distribution of the cell area can be given by

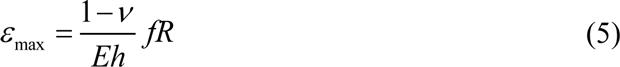

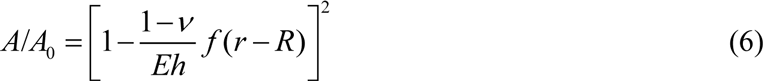

Based on Eq. (4), distribution of the compressive strains in the HeLa cells in experiments can be obtained by analyzing the distribution of cell area, which is shown in Fig. 2b. The sequence in Fig. 2b shows the morphologies of growing HeLa cells with the distribution of the compressive strains given in Fig. 5. The size of the cell monolayer sheet grows bigger along with the cell proliferation, generating higher level of compressive stress in the monolayer. Areas of individual cells in the central region are reduced due to the increased compressive strains. The experimental observations and calculations are consistent with the theoretical predictions. To validate the theoretical model, finite element simulations are performed to explore the relation between the cell-substrate interactions and stress field in growing cells. Results of finite element simulations are shown in Fig. 3d. In the finite element model, the growing cell monolayer sheet was placed on a stiff substrate with the interfacial fraction factor being controlled. The cell sheet was under isotropic expansion to simulate the growth. More than 12,000 linear hexahedral elements were adopted to in the simulations. The cell sheet was modeled as the linear elastic material. With the increase of the growth strain, compressive stresses in the cell monolayer are generated due to the cell-substrate interactions, and the central region has a higher level of compressive stress. The stress level can be reduced by regulating the cell-substrate interactions (Fig. 3d). Finite element simulations are consistent with experimental observation and theoretical analysis.

The elastic strain energy stored in the cell would also increase with the cell proliferation. When the elastic strain energy in the cell is small, interfacial normal adhesion would impose restriction on cell extrusion, making the cell monolayer grow in plane. Thus, higher compressive stress would be generated further. When the compressive stress reaches a critical value, elastic strain energy stored in the cell may be greater than the energy needed for the occurrence of the interfacial delamination. In this critical condition, cell extrusion may happen, and the elastic strain energy can be released. The critical condition for the cell extrusion can be written as

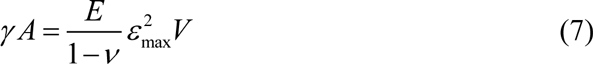

The left hand and right hand of the equation refer to the energy for the interfacial delamination and elastic strain energy in the cell, respectively. *γ* is the energy per area for the interfacial delamination, which is related to the interfacial normal adhesion between the cell and substrate. ε_max_ is the maximum strain in the cell, and *V* is the cell volume. Eq. (6) demonstrates that there exists a critical area or size of the cell monolayer at the critical condition of cell extrusion. The critical size of the cell monolayer sheet depends on the mechanical properties of cell and cell-substrate interactions. The cell extrusion can be controlled by regulating the modulus of the cell and cell-substrate interactions.

When the normal adhesion between the cell monolayer and the substrate is enhanced, cell extrusion would be more difficult to happen, and the cell would bear larger compression before extrusion. When the tangential adhesion is larger, higher level of compression would be induced, and the cell extrusion is more likely to happen. The adhesion between the cells may have effect on the cell sheet stress. Maximum compressive stress in the cell monolayer may be reduced when lowering down the adhesion between the cells. Thus, cell extrusion would be more difficult to happen. Cytoskeleton is the main components that determine the cell mechanics. The cytoskeleton can bear compressive stress and store elastic strain energy. If the cytoskeleton in the cell is suppressed, cell extrusion is hard to happen. These predictions were confirmed by experiments (Fig. 3e-g).

Shape of the cell can be changed due to the compression induced by the interfacial shear stress. Thus, the orientation of the cell division may be altered. During the evolution of the cell monolayer sheet, cell shape in the central region was changed from flat to columnar due to the high level of compression stress. While in the periphery of the monolayer, cell shape maintained flat. Stretch ratio of the cell in the direction of thickness can be written as

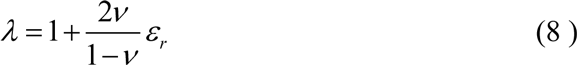

Base on the stretch ratio in Eq. (8), cell thickness/width characterizing cell shape can be obtained as

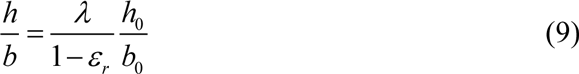

Here *h* and *b* refer to the cell thickness and width. *h*_0_ and *b*_0_ refer to the initialthickness and width of the cell without sustaining compression. In the periphery of the monolayer, cell thickness/width is small since the compressive strain in the cell is small. While in the central region with high level of compressive strain, cell thickness/width can be very large. This phenomenon of cell shape variation induced by the interfacial mechanical interaction is also confirmed by the finite element simulations, as shown in Fig. 4c.

## Supplementary Figures

**Figure S1.**
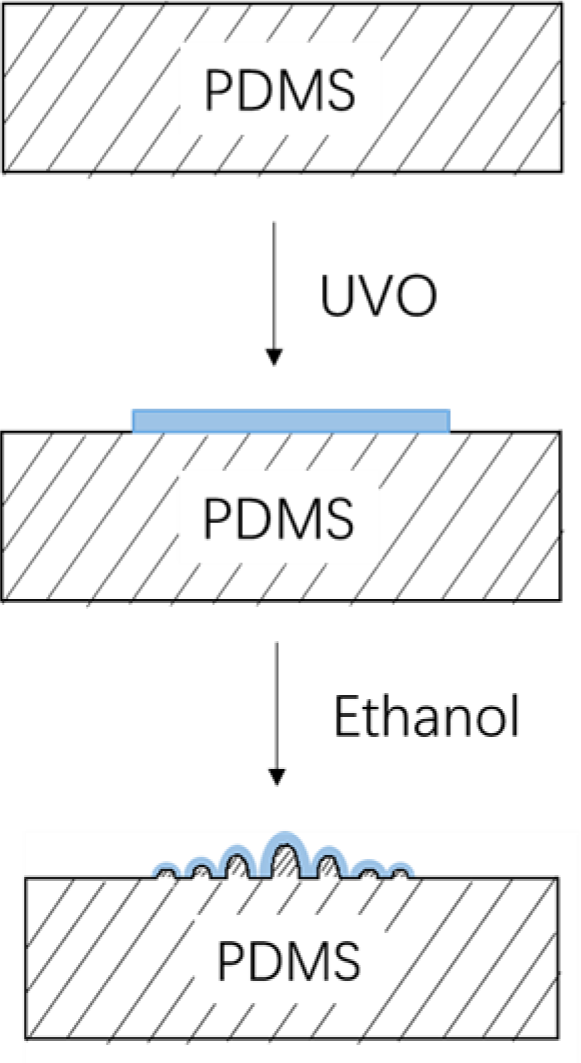
Surface wrinkling induced by differential expansion in a film-substrate system (Zhao et al., 2015b). Polydimethylsiloxane (PDMS) was exposed to UV/Ozone (UVO) for 10 – 55 minutes to form a stiff solvent-responsive oxide layer (highlighted in blue) on the surface. Then dropping an ethanol/glycerol mixture solution containing 60% – 100% ethanol by volume to induce surface wrinkling.

**Figure S2.**
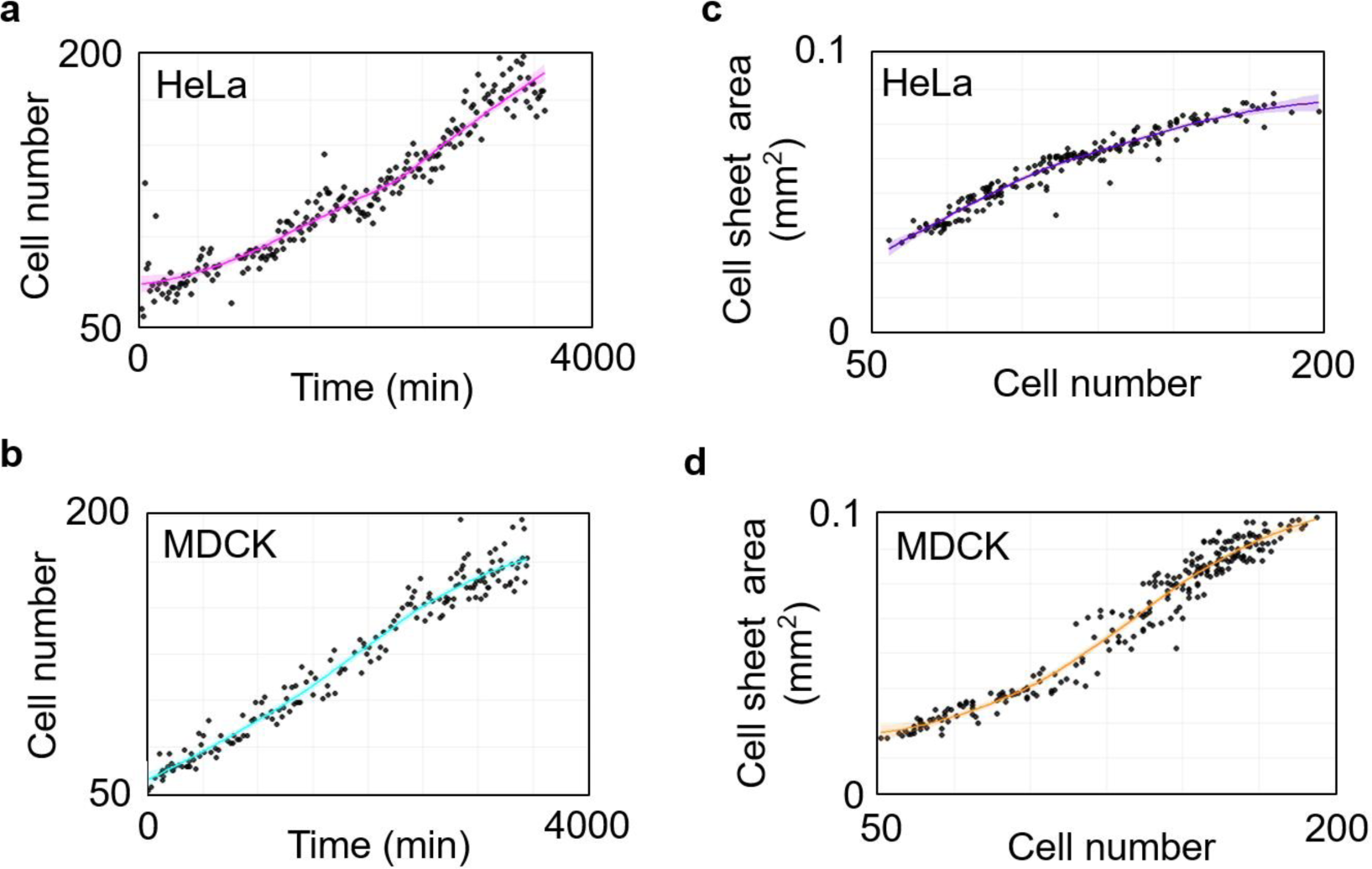
Proliferation and expansion of cell sheets during growth. The statistical analysis of the cell number during HeLa cell sheet (a) and MDCK cell sheet (b) growth. The statistical analysis of the cell sheet area during HeLa cell sheet (c) and MDCK cell sheet (d) growth.

**Figure S3.**
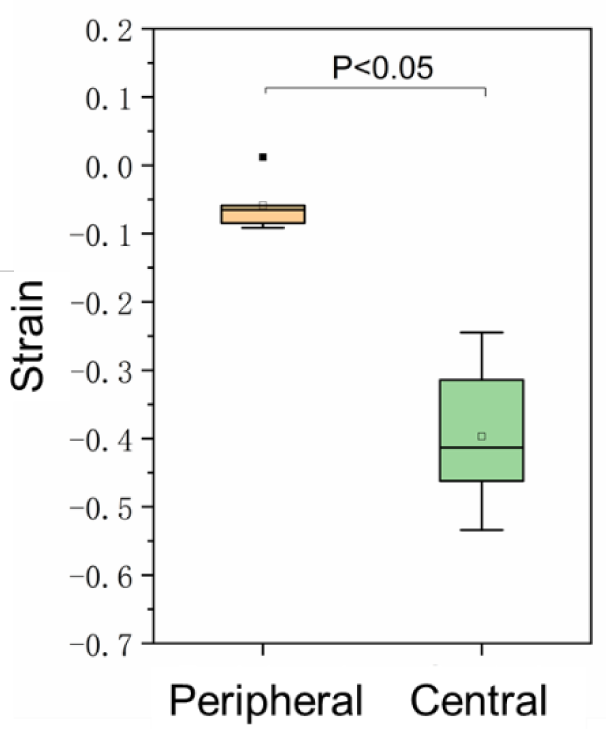
Single cell tracing of compressive strain. The plane strain of single cells in central and peripheral regions of cell sheets during 24 hours analyzed by holographic imaging cytometer (n = 6).

**Figure S4.**
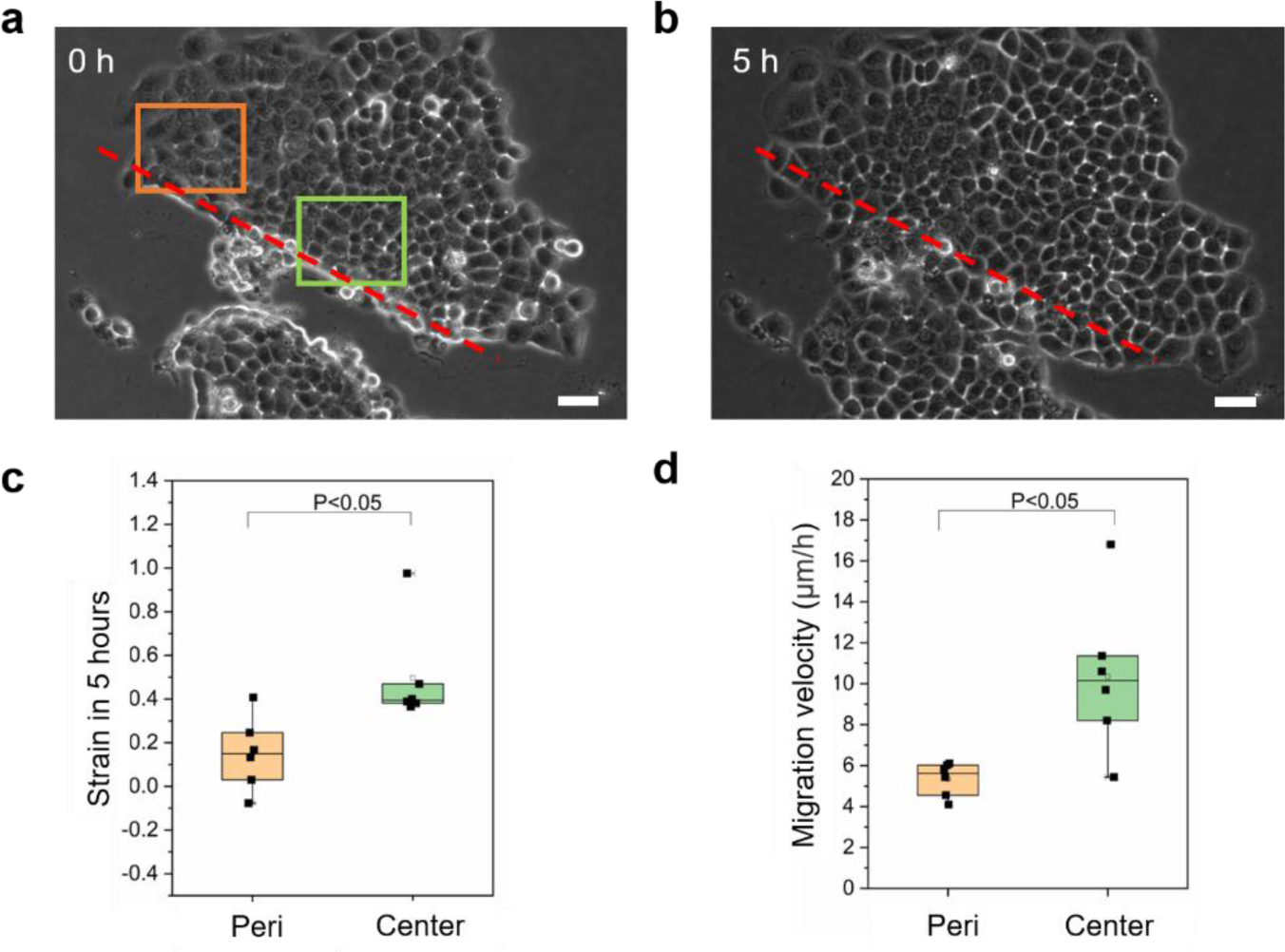
Cell sheet scratch assay. The representative image of cell sheet upon (a) and 5 hours after (b) scratch passing through the center (green box) and edge (orange box) of the cell sheet. The expansion (c) and migration speed (d) were compared between cells in the central region and peripheral region of the cell sheet. Scale bar: 50 μm, (n = 6).

**Figure S5.**
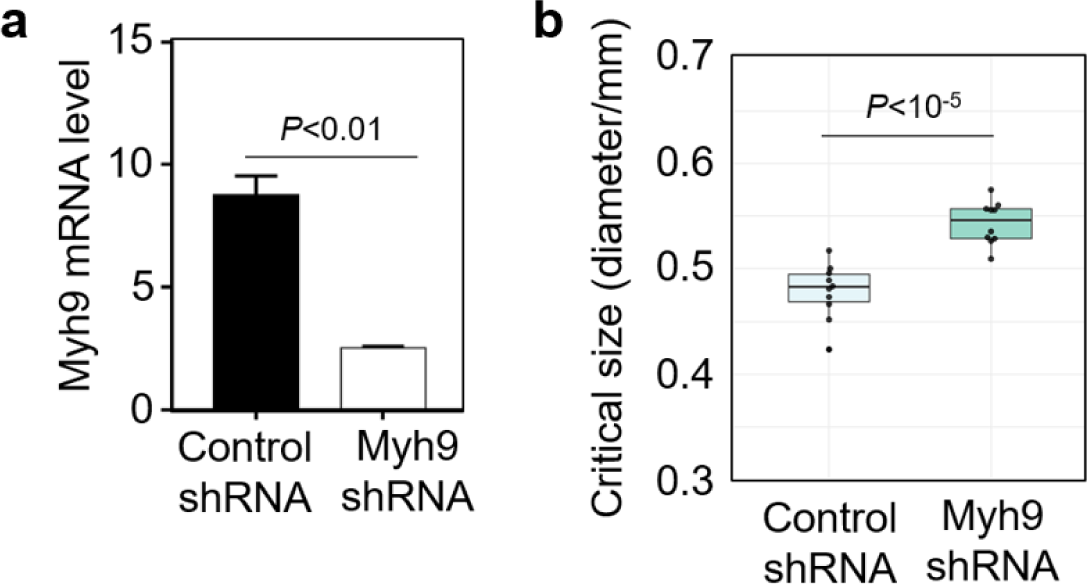
Inhibition of Myosin effects the emergence of 3D morphogenesis. (a) The mRNA level of myosin heavy chain 9 gene (myh9) in HeLa cells transfected with myh9 shRNA and control shRNA. (n = 3). (b) The critical sizes for morphogenesis in myh9 shRNA cells and control cells. (n = 10).

**Figure S6.**
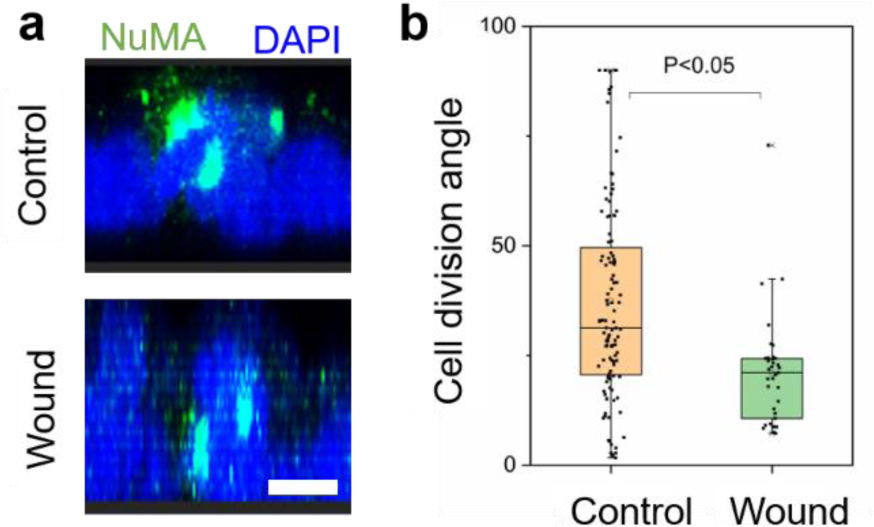
Cell division angle after scratch. (a) Representative XZ slice image of NuMA and nucleus stained by NuMA antibody and DAPI respectively in the control and wound of HeLa cell sheet. (b) The cell division angle in the center of cell sheet without (Control) or with (Wound) scratch. Scale bar: 10 μm.

**Figure S7.**
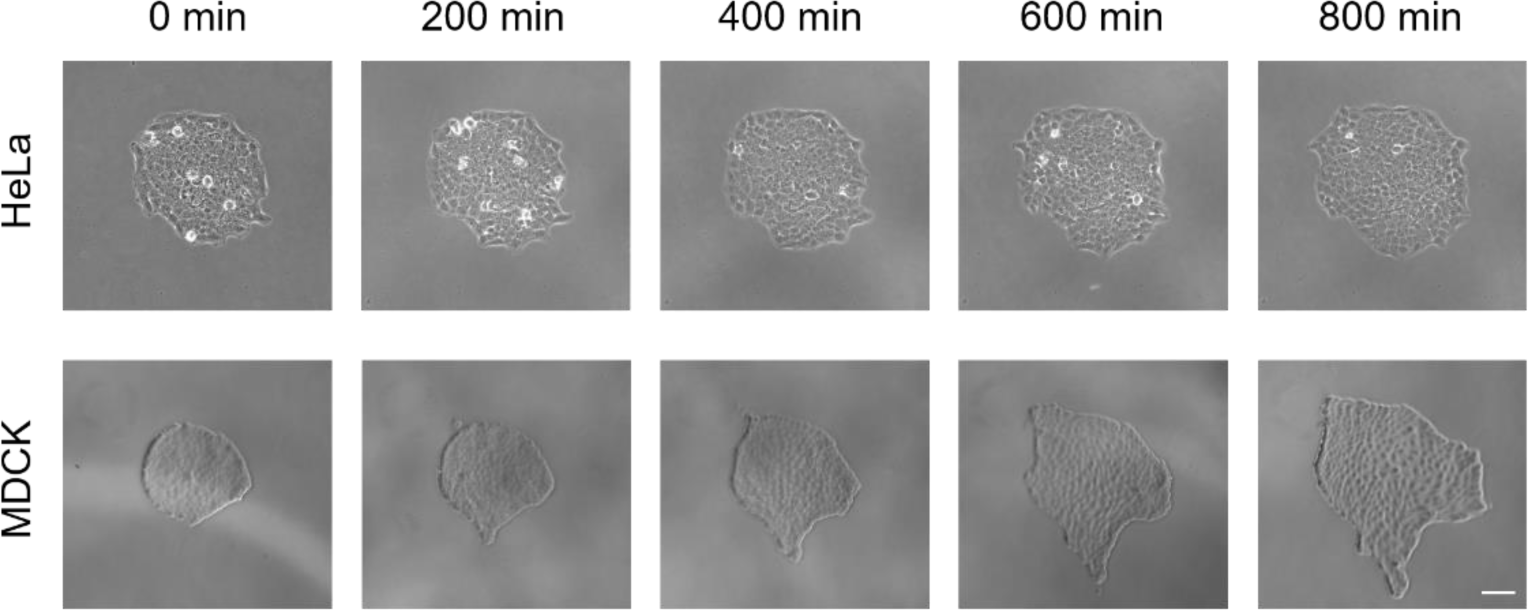
Different deformation pattern between HeLa and MDCK cell sheets during growth. Representative phase contrast images of a growing monoclonal HeLa and MDCK cell sheet captured at the indicated time (min: minute). Scale bar: 100 μm.

**Figure S8.**
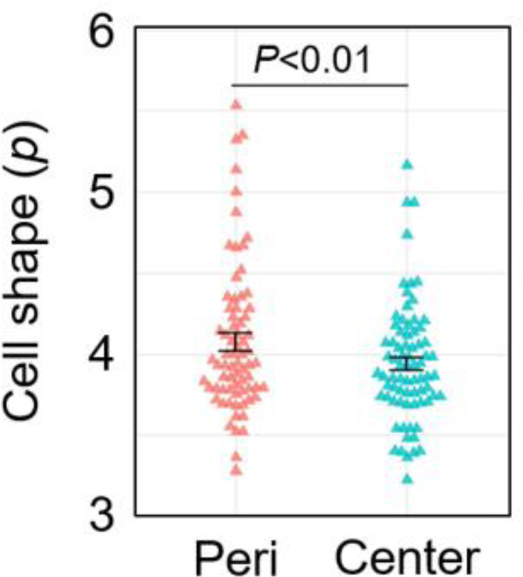
Cell shape index in the peripheral and central regions of HeLa cell sheet. The statistical analysis of cell shape index of cells in the peripheral region and central region of HeLa cell sheet. (n = 75).

**Figure S9.**
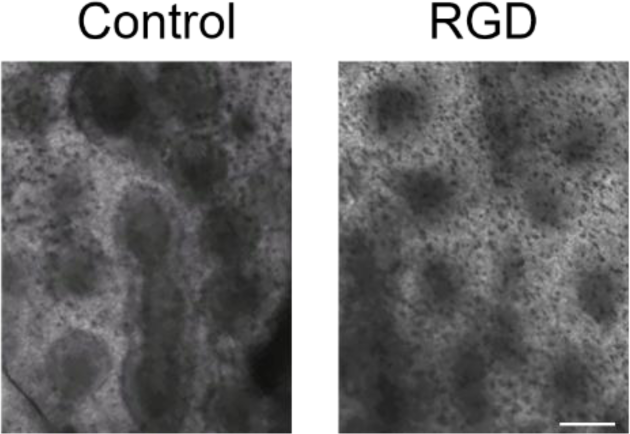
Disrupting the interfacial mechanical interaction between epidermal and dermal layers attenuated the morphogenesis of primordium. *Ex vivo* culture of embryonic chicken skin with or without RGD. Scale bar: 500 μm.

## Supplementary Table

**Table S1.** The critical size for 3D morphogenesis of growing cell sheet on different types of ECM and stiffness (polyacrylamide gel).

| ECM type | Critical diameter($\mu\text{m}$) | N |
| --- | --- | --- |
| No pre-coating ECM | 463.8±18.9 | 10 |
| Fibronectin | 493.8±23.9 | 10 |
| Gelatin | 489.1±19.8 | 10 |
| Collagen | 520.3±21.8 | 10 |
| Collogen (40 kPa PA gel) | 413.8±9.6 | 5 |
| Collogen (1 kPa PA gel) | 342.6±26.0 | 4 |

## Supplementary Movies

**Supplementary Movie 1.** Symmetric cell division before critical compression during HeLa cell sheet growth visualized by HoloMonitor M4 time-lapse cytometer.

**Supplementary Movie 2.** Asymmetric cell division after critical compression during HeLa cell sheet growth visualized by HoloMonitor M4 time-lapse cytometer.

**Supplementary Movie 3.** PIV analysis during HeLa and MDCK cell sheets growth.

**Supplementary Movie 4.** Time-lapse imaging of MDCK cell sheet during growth.

**Supplementary Movie 5.** PIV analysis during MDCK cell sheet growth in the presence of inhibitors or vehicle (control).

